# Maternal nutritional stress alters sperm competence in male mice offspring leading to reduced fertility

**DOI:** 10.1101/2020.11.10.376616

**Authors:** Yuki Muranishi, Laurent Parry, Mélanie Vachette-Dit-Martin, Fabrice Saez, Cécile Coudy-Gandilhon, Pierre Sauvanet, David Volle, Jérémy Tournayre, Serge Bottari, Francesca Carpentiero, Jessica Escoffier, Alain Bruhat, Anne-Catherine Maurin, Julien Averous, Christophe Arnoult, Pierre Fafournoux, Céline Jousse

## Abstract

Infertility represents a growing burden worldwide, with one in seven couples presenting difficulties conceiving. Amongst these, 10-15% of the men have idiopathic infertility that does not correlate with any defect in the classical sperm parameters measured. In the present study, we used a mouse model to investigate the effects of maternal undernutrition on fertility in male progeny. Our results indicate that mothers fed on a low protein diet during gestation and lactation produce male offspring with normal sperm morphology, concentration and motility but exhibiting an overall decrease of fertility when they reach adulthood. Particularly, sperm from these offspring show a remarkable lower capacity to fertilize oocytes when copulation occurs early in the estrus cycle relative to ovulation, due to an altered sperm capacitation.

Our data demonstrate for the first time that maternal nutritional stress can have long-term consequences on the reproductive health of male progeny by affecting sperm physiology, especially capacitation, with no observable impact on spermatogenesis and classical quantitative and qualitative sperm parameters. Moreover, our experimental model could be of major interest to study, explain, and ultimately treat certain categories of infertilities.

## Introduction

The “developmental origins of health and disease” (DOHaD) hypothesis proposes that maternal nutritional status during pregnancy and lactation plays a major role in postnatal growth and development of the offspring, determining long-term health status. This process, termed nutritional programming, allows specific adaptations to be made that permanently change the phenotype of the organism. These changes persist even in the absence of the early nutritional stress that initiated them, leading to lifelong health consequences. In humans, a considerable amount of evidence indicates that maternal malnutrition during gestation can confer an increased risk of metabolic dysfunction in offspring^1 2^. In mice, several studies have also shown that maternal nutritional programming has long lasting effect on offspring metabolism^3,4^. For instance, we have demonstrated that pups born from mothers fed a Low Protein Diet (LPD) during gestation and lactation have an altered metabolic status in adulthood, displaying lower body weight, lower fat mass and hypermetabolism^5,6^.

However, very few studies have explored the effect of nutritional programming on other physiological functions, such as reproductive function. Infertility is increasing worldwide, and it currently affects about one in seven couples attempting to conceive^7^. In about 50% of cases, infertility is due to the male partner^8^. Meta-analyses^9^ indicate that there has been a considerable decline in sperm quality since the 1940s suggesting a defects in spermatogenesis. However, a significant proportion of male infertility is idiopathic and not related to defect in spermatogenesis. Whatever the case, the reason for this infertility is not well understood, and it probably has multifactorial causes. Some authors suggest that reproductive disorders may have a developmental origin^10^. Indeed, a few studies report that, in sheep and rodents, maternal dietary restriction impairs reproductive performance in female offspring^11,12^. Similarly, testicular development and reproductive performance of male offspring has been linked to maternal protein restriction^13^. However, the underlying causes are unknown.

Here, we investigated the effects of maternal undernutrition on the offspring’s fertility. We used the well-characterized model of dams fed an isocaloric LPD during both gestation and lactation. Our results indicate that low dietary protein intake by dams is associated with reduced fertility in male offspring due to altered sperm fitness. For these males, fertilization capacity is reduced when copulation takes place early in the estrus cycle relative to ovulation.

## Results

### Maternal undernutrition alters fertility in offspring

Female Balb/c mice were fed either a control diet (CD) or an LPD throughout pregnancy and lactation. After weaning, the F1-CD and F1-LPD offspring – born and suckled from CD- or LPD-fed mothers, respectively – were fed a standard chow diet. At the age of six months, both male and female F1-LPD exhibited a lower body weight than their F1-CD counterparts (Figure 1A). Moreover, male F1-LPD offspring had a significantly lower perigonadal-White Adipose Tissue weight (pg-WAT). Even though a similar pg-WAT difference was observed in F1-LPD females, due to greater interindividual variability in females the difference was not significant (Figure 1A). This gross anatomical phenotype was also associated with a shorter AnoGenital Distance (AGD) in F1-LPD males compared to F1-CD males. AGD is a marker of differentiation of male and female external genitalia, and a lower AGD is commonly used as a predictive indicator of diminished reproductive function in the mature male^14,15^. To adjust AGD to compensate for differences in weight between animals, Gallavan et al.^16^ reported that the cube root of body weight would be an appropriate standard to use. The corrected factor, called AGI (anogenital Index), was also significantly reduced in F1-LPD males compared to F1-CD males. Interestingly, even though AGD may also have an impact on female fertility^17^, maternal undernutrition affected neither AGD nor AGI in F1-LPD females (Figure 1A).

**Figure 1.**
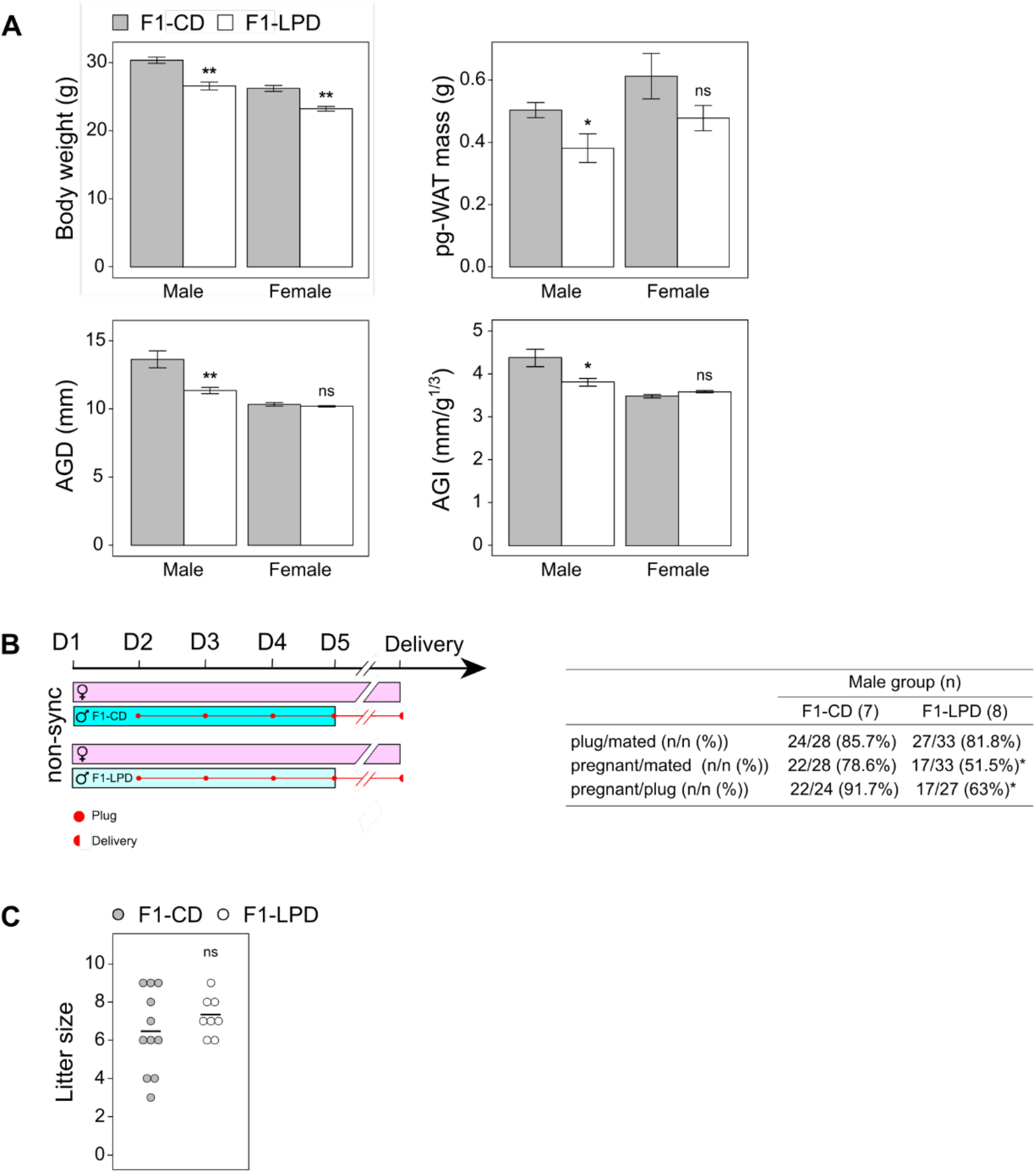
Reproductive characterization of F1 offspring. <u>**(A)** Phenotypic characterization of F1 offspring</u>. Body weight, peri-gonadal White Adipose Tissue mass (pg-WAT), AnoGenital Distance (AGD) and AnoGenital Index (AGI) are given for F1-CD and F1-LPD adult males and females at 6 months of age. AGI is defined as AGD/(BW)^1/3^. Results are expressed as means +/- SEM for at least 6 individuals per group and student’s t-test was used to test the significance (** p-value < 0.01, * p-value < 0.05, ns= non-significant). For non-invasive measures (BW, AGD and AGI) similar data obtained from a large number of individuals have not been included in the figure. <u>**(B)** Fertility evaluation of F1-CD and F1-LPD males when mated with Balb/c females</u>. F1-CD and F1-LPD males (blue bars) were caged with non-synchronized Balb/c females (pink bars) at Day 1 (D1) at a 1:2 male to female ratio for 4 days (until D5). Vaginal plugs were checked every day (●) and plugged females were removed and isolated until delivery 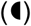. For each group, the number of males tested, total females, plugged females and pregnant females were indicated in the table. Fertility rates are calculated as the percentage of plugged females that are pregnant. Statistical significance was calculated using a Khi-squared test. * indicates a p-value < 0.05, ns= non-significant. <u>**(C)** Litter size of offspring obtained after mating of F1-CD and F1-LPD males with non-synchronized Balb/c females</u>. Each dot represents the litter size of offspring born to 1 Balb/c Female mated with either F1-CD or F1-LPD males. The black horizontal line represents the mean per group and student’s t-test was used to test the significance (ns= non-significant).

In light of the observation of indicators of potential fertility problems, we measured fertility rates for both F1-CD and F1-LPD animals. First, adult F1-CD or F1-LPD males were mated with control females. We observed no difference in the number of plugged females relative to the total number of mated females between the two groups (86% with F1-CD vs 82% with F1-LPD males). Thus, copulation occurred at the same rate for F1-LPD and F1-CD males. However, the fertility rate recorded (Figure 1B) – calculated as the percentage of plugged females that went on to develop a pregnancy – was significantly different between the two groups. Reduced fertility was noted for F1-LPD males (63%) compared to F1-CD males (92%). Thus, the fertility of male mice is affected by maternal perinatal undernutrition. Interestingly, litter size was not significantly different between the two groups (Figure 1C). Based on these results, we can conclude that the reduced fertility observed for the F1-LPD males was thus not the result of behavioral problems, such as prostration, or of production of completely fertilization-incompetent sperm.

In contrast, when F1-CD or F1-LPD females were mated with control males, fertility in F1-LPD females was only slightly lower than in F1-CD females, and the difference was not statistically significant (Figure S1). Maternal nutritional stress may affect fertility in female offspring through several mechanisms. Firstly, it can affect gonad development (even though the female AGD was not significantly altered). Secondly, it is also possible that the maternal LPD diet causes several metabolic/endocrine alterations that could create an inadequate metabolic environment for optimal pregnancy. To definitively determine whether female fertility is affected by maternal nutritional stress, further experiments would be required which are out of the scope of this study. For the remainder of this study, we decided to focus our investigations on the causes of the decline in male fertility.

### Maternal protein malnutrition effects on sexual organ development and sperm count

It is considered useful to measure the mass of sexual organs such as the testis, epididymis, and seminal vesicle to assess alterations to the male reproductive system. We thus first investigated whether maternal undernutrition affected the male reproductive organs in adult mice. Figure 2A shows that the masses of testes and epididymis from adult F1-LPD mice were significantly lower than those from control mice. However, the weight of the seminal vesicles did not appear to be strongly affected. In normal conditions, it is generally accepted that testes weight varies little between individuals from the same species^18^. However, the significantly lower testis weight recorded for F1-LPD mice compared to F1-CD could be linked to the lower body weight observed.

**Figure 2.**
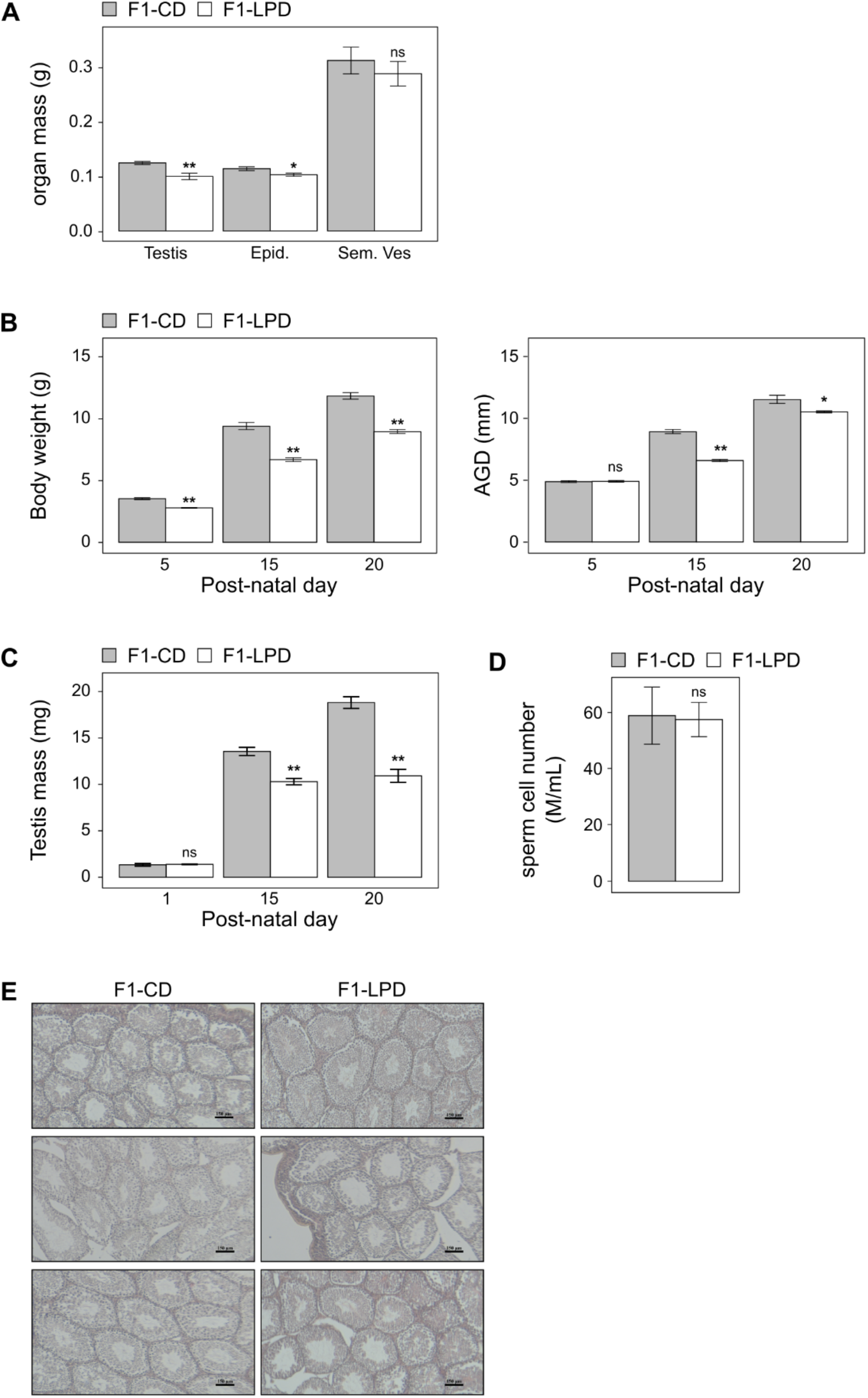
Reproductive organs analysis. <u>**(A)** Reproductive organ masses of F1 adult males</u>. Testis, Epididymis (Epid.) and Seminal Vesicles (Sem.Ves) masses are given for F1-CD and F1-LPD males at 6 months of age. Results are expressed as means +/- SEM for at least 6 individuals per group and student’s t-test was used to test the significance (** p-value < 0.01, * p-value < 0.05, ns= non-significant). <u>**(B)** Body weight (g) and AnoGenital Distance (AGD, mm) of F1-CD and F1-LPD males during suckling period</u>. Body weight and AGD are given for F1-CD and F1-LPD males at 5, 15 and 20 post-natal day. Results are expressed as means +/- SEM for at least 6 individuals per group and student’s t-test was used to test the significance (** p-value < 0.01, * p-value < 0.05, ns= non-significant). <u>**(C)** Testis weight of F1-CD and F1-LPD males during suckling period</u>. Testis weight are given for F1-CD and F1-LPD males at 1, 15 and 20 post-natal day. Results are expressed as means +/- SEM for at least 6 individuals per group and student’s t-test was used to test the significance (** p-value < 0.01, ns= non-significant). <u>**(D)** Sperm count</u>. Number of sperm cells following isolation from the cauda epididymis of 2-month old mice are expressed as millions of sperm cells per mL (M/mL) Results are expressed as means +/- SEM for at least 4 individuals per group and student’s t-test was used to test the significance ns= non-significant). <u>**(E)** Hematoxylin/Eosin (H&E) staining of testis from 2month old mice</u>. (n=3 for each group) Scale bar = 50μm

To determine whether this was the case, we calculated the testis/body weight ratio for Balb/c mice. Figure S2 shows that, across a large group of mice raised in conditions without nutritional stress, there was no correlation between the weight of the animal and the testicle weight. These data suggest that the smaller testis weight recorded for the F1-LPD mice is not due to their reduced overall stature, but is a consequence of the nutritional stress experienced during early life. As a counter example, our data revealed a strong correlation between the animals’ weight and that of their epidydimal adipose tissue.

Based on these findings, we explored the hypothesis that a developmental defect occurred during the period of nutritional stress. In males, testicular cell lines differentiate, proliferate, and mature during postnatal life, particularly during suckling. These processes are crucial in establishing future reproductive capacity. Stresses experienced during suckling could thus impair fertility in adult animals^19^. Figure 2B shows that body weight was significantly lower in F1-LPD animals from 5 days after birth. However, significant differences in AGD (Figure 2B) and testis weight (Figure 2C) were only observed after 15 days. These observations suggest that the lactation period could be critical in determining the impact of maternal nutrition on the future reproductive capacity of male offspring. Despite the anatomical differences recorded for F1-LPD testis, spermatogenesis does not appear to be altered. Indeed, sperm collected in the cauda epididymis had similar cell morphology (not shown) and were present in equivalent numbers (Figure 2D) between F1-CD and F1-LPD males. These results were reinforced by histological analysis of testis sections, showing no differences between the two groups of mice (Figure 2E).

### Maternal protein malnutrition reduces the duration of sperm fertility in the female reproductive tract

As mentioned above, the reduced fertility observed for the F1-LPD males seemed to be neither the result of a behavioral problem, nor of a problem with sperm production. Thus, it is reasonable to hypothesize that sperm fitness is affected in these animals. For a given species, in normal conditions, the lifespan of sperm cells in the female reproductive tract is adapted to the interval between the onset of estrus and ovulation, allowing sperm to survive in a fertilization-competent form until ovulation occurs^20^. However, in rodents, there is a certain disparity concerning the timing of copulation relative to the female’s estrogen cycle. In mice (as it is the case in human), males mount females regardless of their estrus state^21^. We therefore considered whether the timing of copulation, relative to the female’s estrus cycle, could be a determinant factor in the reduced fertility observed for F1-LPD males. We speculated that the fertilizing capacity of the F1-LPD male semen might be affected by the interval between copulation and ovulation. To test the hypothesis that longevity and/or fertilizing capacity of sperm cells in the female genital tract might differ between F1-CD and F1-LPD males, we used a well-established protocol to synchronize ovulation in control females by exposure to gonadotropins.

In a first experimental procedure (Figure 3, protocol A), males were mated with females overnight (i.e. over a short time window) to allow copulation only at the time of the hormonally-stimulated ovulation (ovulation is indicated in Figure 3 by a red lightning bolt). In this set-up, no differences in the rate of female pregnancy were observed between F1-CD and F1-LPD males, thus sperm produced by F1-LPD males could fertilize oocytes. In contrast, in a second experimental procedure (Figure 3, protocol B), males were mated with females in a copulation window of 57 h to 42 h before ovulation (considering ovulation to occur at T0). In these conditions, the F1-CD males could still successfully fertilize the females, but the capacity of the F1-LPD males to do so was severely compromised. These data indicate that F1-LPD sperm could no longer fertilize oocytes if they remained in the female reproductive tract for too long before ovulation. This result suggests that the fertilizing capacity of the sperm cells produced by F1-LPD males decreased rapidly once in the female reproductive tract.

**Figure 3.**
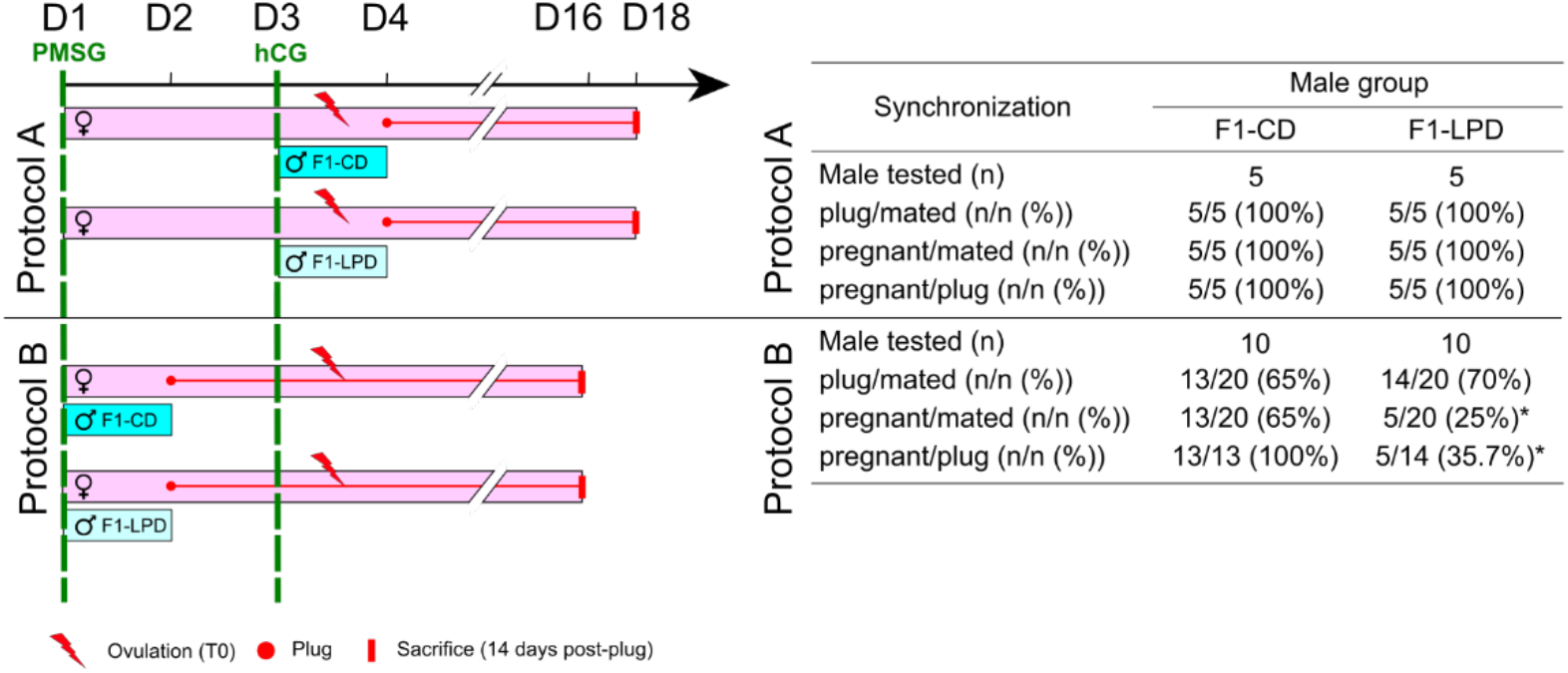
Fertilizing capacity evaluation of F1-CD and F1-LPD males mated with synchronized Balb/c females. F1-CD and F1-LPD males (blue bars) were caged with Balb/c females (pink bars) at a 1:1 (Protocol A) or 1:2 (Protocol B) male to female ratio for 1 day according to 2 different protocols. In both cases, females were hyper-ovulated and synchronized by intraperitoneal injection of PMSG (at D1, green dashed line) followed by hCG 48h later (at D3, green dashed line). Ovulation takes place the night following hCG injection, i.e. between D3 and D4, characterized by a red lightning bolt. In protocol A, males and females were caged together during the ovulation period, between D3 and D4. In protocol B, males and females are caged together between D1 and D2, about 48h before ovulation. For both protocols, vaginal plugs were checked (●) at the end of the copulatory period and plugged females were removed and isolated for 14 days. Females were thus sacrificed 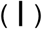 in order to evaluate pregnancy. For each group and each protocol, the number of males tested, total females, plugged females and pregnant females were indicated in the table. Fertility rates are calculated as the percentage of plugged females that are pregnant. Statistical significance was calculated using a Khi-squared test. * indicates a p-value < 0.05.

### Maternal protein malnutrition alters capacitation process

Sperm are maintained inactive in the epididymis for relatively long periods of time (1–2 weeks, depending on the species). When sperm are released into the female reproductive tract, the change in environment acts as a physiological trigger, and sperm start their capacitation process. This process, necessary to allow oocyte fertilization^22,23^, involves remodeling of lipids and proteins^24^, resulting in alterations to membrane fluidity^25^. Capacitation is also associated with an increase in oxidation levels that may affect the sperm’s lifespan (DNA and lipid damage lead to a loss of sperm motility and a commitment to cell death)^26,27^.

To understand the underlying causes of the observed alteration in fitness of sperm produced by F1-LPD males, we first used a Computer-assisted Semen Analyzer (CASA) to assess physiological parameters in sperm isolated from the cauda epididymis. As shown in Figure 4A, after 1 h of *in vitro* capacitation, the percentage of motile F1-LPD sperm cells was significantly decreased compared to F1-CD samples whereas no such difference was observed before capacitation. Moreover, the percentage of hyperactivated sperm cells after 1 h of *in vitro* capacitation was also significantly decreased in F1-LPD compared to F1-CD. These data led us to conclude that, before the start of capacitation, sperm cell motility is no different between the two groups, but that differences appear in response to capacitation signals. From these observations, we propose two non-exclusive hypotheses: the *in vitro* capacitation process is (i) either less efficient in F1-LPD sperm compared to F1-CD and/or (ii) leads to a premature commitment of F1-LPD sperm to cell death.

**Figure 4.**
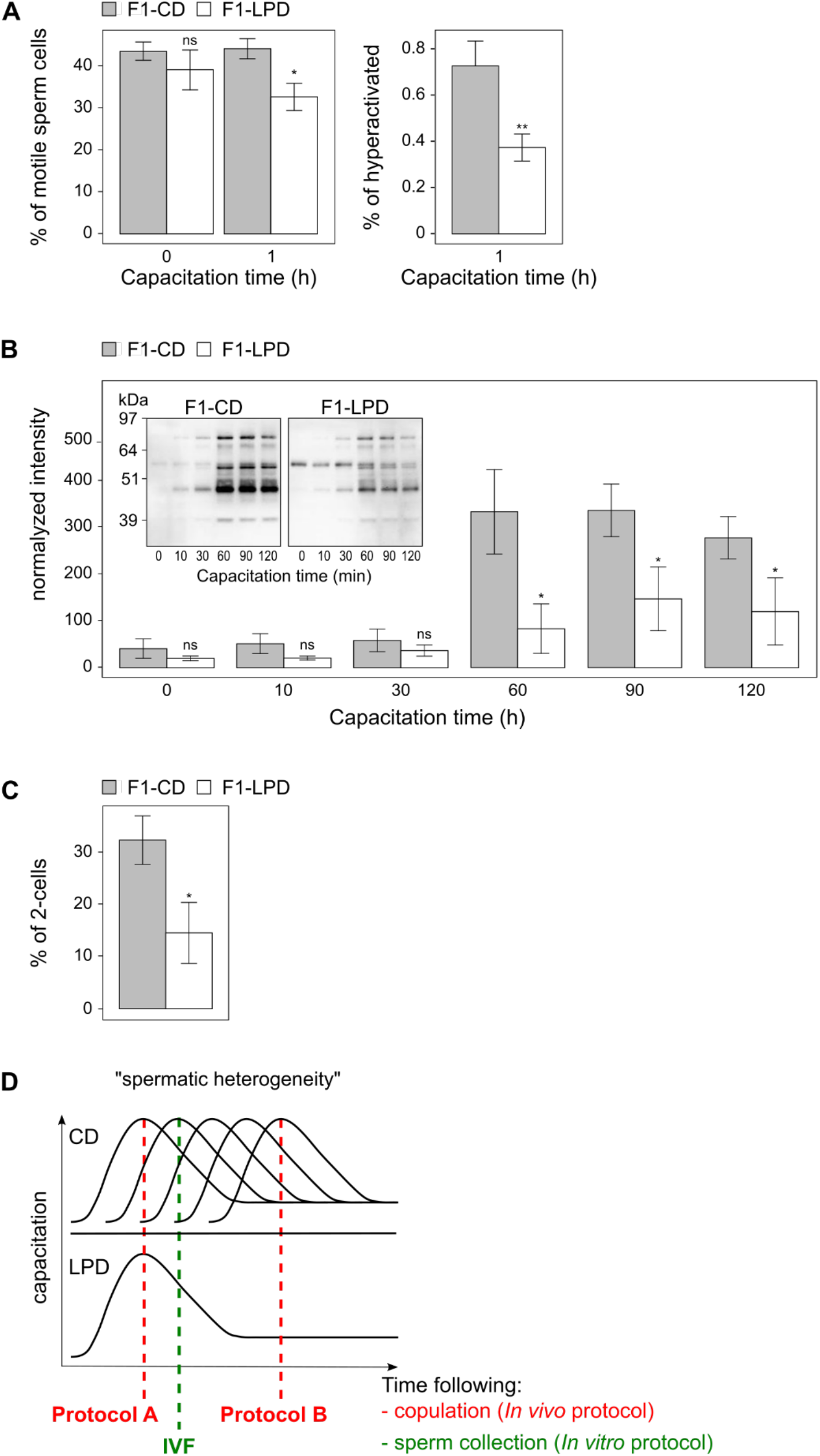
Sperm cell parameters and functionality. <u>**(A)** Sperm cell parameters (motility and activation)</u>. Motility and activation parameters (% of hyperactivated) evaluated by CASA (Computer Aided Sperm Analysis) are shown for sperm cells before capacitation and after 1h of capacitation. Mean +/- SEM for each parameter is shown for n=8 in the F1-CD group and n=11 for the F1-LPD group. Student’s t-test was used for comparison between groups. Statistical significance is indicated on the graphs: * p-value < 0.05, ** p-value < 0.01, ns= non-significant). <u>**(B)** Capacitation efficiency determined with anti-phosphotyrosine Western blot</u>. Identical numbers of sperm cells, isolated from cauda epididymis of 3 F1-CD and 3 F1-LPD mice, were incubated 0, 10, 30, 60, 90, 120 min in classical capacitation medium for *in vitro* capacitation kinetics. Protein extracts were analyzed by Western bloting analysis of protein Tyrosine phosphorylation using anti-phosphotyrosine antibody. Protein loads were controlled with TGX stain free precast gels and a quantification was realized using ImageJ software. Mean +/- SEM is shown for n=3 in each group. Student’s t-test was used for comparison between groups. Statistical significance (*) is indicated on the graphs for p-value <0.08. The blot given is representative of semen from three separate individuals of each group. <u>**(C)** Fertilization competence of sperm isolated from F1-CD and F1-LPD males using in vitro fertilization assay</u>. Sperm were isolated from 4 males of each group and incubated in capacitating medium for 12h. Oocytes were isolated from Balb/c hyperovulated females. Fertilization competence was measured by determining the percentage of two-cell embryos obtained 24h after fertilization of at least 10 oocytes per male sample. <u>**(D)** Schematic representation of the spermatic heterogeneity hypothesis</u>.

To explore further the impact on capacitation, we measured and analyzed the level of protein tyrosine phosphorylation (the gold standard for assessing capacitation) from F1-CD and F1-LPD sperm. Data given Figure 4B show a quantification of three separate individuals per group as well as a representative blot. It shows that the intensity of the bands representing tyrosine-phosphorylated proteins in F1-CD sperm extracts was maximal after 60 min incubation in capacitating medium. Interestingly, although the kinetic profile for tyrosine phosphorylation was no different for F1-LPD sperm extracts, the phosphorylation intensity was significantly lower after 60 min. These results indicate that the capacitation process was strongly affected in F1-LPD sperm cells.

Capacitation is essential to successful oocyte fertilization, and any defects in this process compromise the fertilization competence of sperm. Overall, our results strongly suggest that F1-LPD sperm are less fit, thus their fertility may be reduced in challenging conditions. To test this hypothesis, we determined how many two-cell embryos were obtained upon *In Vitro* Fertilization (IVF) of oocytes with either F1-CD or F1-LPD sperm. In these experiments, the percentage of 2-cell embryos is regarded as the *in vitro* fertilization rate. We chose to incubate sperm in a capacitation medium for a long challenging time (12 h) before mixing gametes as a simulation of what occurs in the genital female tract *in vivo* when copulation takes place a long time before ovulation. Figure 4C shows that after 12 h of *in vitro* capacitation, the *in vitro* fertilization rate was strongly reduced for F1-LPD sperm compared to F1-CD sperm. Thus, after a long *in vitro* capacitation step, F1-LPD sperm were less capable of fertilizing oocytes.

Overall, these data show that a low maternal dietary protein intake during gestation and lactation is associated with altered sperm fitness in the offspring, resulting in a lower capacity to successfully fertilize oocytes when copulation takes place early before ovulation.

## Discussion

The aim of this article was to determine whether a maternal diet low in protein affected fertility in offspring at adulthood. Our results indicate that maternal nutritional stress during gestation and lactation decreased fertility in male offspring.

### 1/ Sperm capacitation is altered in F1-LPD sperm

Our results demonstrate that sperm fertilizing competence is altered due to an important capacitation defect. The capacitation process corresponds to the various changes that are required for sperm cells to become capable of fertilizing oocytes. It occurs as sperm pass through the female reproductive tract. As sperm are transcriptionally and translationally inactive, all the changes leading to capacitation are controlled by complex signaling cascades involving post-translational modifications of proteins (phosphorylations) and redox processes^28^. Thus, sperm capacitation involves a series of primary events such as influx of extracellular Ca^2+^ and Na^+^, increase in cyclic-AMP, decrease in intracellular pH, changes in membrane phospholipid content and loss of membrane cholesterol, leading to alterations to membrane fluidity^25^. In sperm from F1-LPD males, both the overall intensity of tyrosine phosphorylation (a marker of capacitation), and the proportion of hyperactivated sperm cells following *in vitro* capacitation were decreased compared to control sperm. These observations suggest a defect in the sperm’s response to the capacitation signal, which may be caused by several factors, acting in isolation or in combination.

#### Deregulation of the redox balance

Whereas physiological levels of ROS are vital for optimal sperm function, when present at excess levels, ROS adversely affect sperm quality and function, ultimately resulting in infertility^26,27^. Interestingly, mitochondrial function is affected in animals born to mothers fed an LPD during gestation and lactation. Indeed, our previous studies indicated that F1-LPD animals present increased mitochondrial function in skeletal muscle and have an increased mitochondrial density in White Adipose Tissue^5^. Whether this increase in mitochondrial function also trigger oxidative stress in sperm cells has yet to be addressed. However, gene expression data (not shown) indicate that antioxidant enzymes such as GPX1, GPX3 and SOD3 are increased (1.3-fold each) in the epididymis in F1-LPD animals. This increased expression could be due to a defensive response to oxidative stress. As mentioned above, oxidative stress and excessive ROS production may affect sperm parameters such as motility (see^29,30^ for review). The altered motility observed in F1-LPD sperm during *in vitro* capacitation (Figure 4A) might be the result of excess ROS, and thus increased oxidative stress.

#### Role of leptin

Leptin is a well-characterized metabolic actor involved in the regulation of fatty acid and cholesterol metabolism, which may play important roles in male reproduction. First, leptin plays a crucial role at the onset of puberty. For example, leptin signaling may be involved in the delayed puberty reported in pups of both sexes born from mothers exposed to perinatal food restriction^31^. In pigs, leptin has also been shown to influence sperm capacitation by enhancing cholesterol efflux and protein tyrosine phosphorylation^32^. Since leptin plasma concentration is strongly decreased in F1-LPD animals^6^, this could affect sperm capacitation through its effect on lipid metabolism.

### 2/ Fertile lifespan of F1-LPD sperm is compromised

For a given species, the fertile lifespan of sperm correlates directly with the interval between the onset of estrus and ovulation. In normal conditions, successful fertilization relies on heterogeneity among sperm cells. Since capacitation is a time-limited process, various sperm subpopulations co-exist and can capacitate at different times^33,34^ (Figure 4D). This particularity responds to a physiological need to ensure that sperm will remain fertilization-competent until ovulation takes place. We have clearly shown that F1-CD males can fertilize females *in vivo*, whenever copulation occurs relative to ovulation. In contrast, sperm from F1-LPD males were unable to induce fertilization if copulation was not timed to coincide with ovulation. Similarly, IVF experiments showed that the percentage of sperm cells that lose their fertilizing competence following a long capacitation (12 h) is higher for sperm from F1-LPD than from F1-CD.

Altogether, these results demonstrate that maternal nutritional stress affects sperm physiology and particularly the *in vivo* sperm lifespan, but has no effect on spermatogenesis *per se*. The altered sperm lifespan may be due to (i) defective capacitation once in the female reproductive tract and/or (ii) decreased sperm heterogeneity due to a lack of the subpopulations of sperm cells showing delayed capacitation (Figure 4D).

In humans, it is important to note that 10 to 15% of men who consult for difficulties conceiving present an idiopathic infertility, without defects in the classical sperm parameters measured, including sperm morphology and concentration, DNA compaction and DNA breaks and sperm motility^35^. Sperm heterogeneity and sperm physiology are very rarely assessed and in particular sperm capacitation is never measured, and any defects affecting such a parameter could play a particularly important role in idiopathic infertility. In that context, identification of molecular mechanisms involved in the establishment and maintenance of the imprint leading to infertility in our mouse model will allow to study, explain, and ultimately treat these categories of infertilities in humans. Moreover, it would also be interesting to assess whether, in such patient, idiopathic infertility correlates with maternal nutritional stress.

## Materials and Methods

### Ethics statements

Maintenance of the mice and all experiments were approved by our institutional-animal care and use committee in conformance with French and European Union laws (permission to experiment on mice B63-150, local ethic committee CEMEAA18-13, animal facilities agreement C6334514, project agreement APAFIS#13710-2016122214418591 v7).

### Experimental Design

All mice were housed individually in polycarbonate, standard mouse cages (31l*20w*15h cm3) containing wood chips bedding and a plastic tube for enrichment and were changed to clean cages every two weeks. The ventilated housing room was maintained at a 12:12 hour light-dark cycle beginning at 8 a.m., at an ambient temperature of 22±1°C and 60±5% humidity. All mice were fed ad libitum and given free access to water. Experimental diets were manufactured in our institute facilities (INRAE de Jouy-en-Josas, Sciences de l’Animal et de l’Aliment de Jouy–Régimes à Façon, Jouy-en-Josas, France).

Pairs of 10-week-old Balb/c virgin female mice from Janvier Labs (53941 Saint-Berthevin, France) were mated with a single male. At detection of vaginal plugs, females were allocated into 2 groups and fed either a Control Diet (CD) containing 22% protein or a Low Protein Diet (LPD) containing 10% protein throughout gestation and lactation as previously described in^6^. Only litters of 4–10 pups were included in subsequent experiments. After weaning at 4 weeks of age, the F1-CD and F1-LPD offspring, born respectively to CD or LPD-fed mothers were single-housed and were given standard chow diet (A03; Safe, Augy, France) throughout life.

### Body weight and anogenital distance measurement

Body weight and anogenital distance (AGD) of all offspring were measured at 5, 15, 20 days and 6 months of age. Body weight was measured using a standard top-loading laboratory balance. AGD (expressed in mm) was measured with a digital caliper from the center of the anus to posterior edge of the genital papilla, by a single investigator to increase precision. Pups were gently restrained, and the anogenital area was subjected to slight tension to impart tautness to the region AGD measurements were corrected by cubic root of body weight (Anogenital index = AGI= AGD/(BW)^1/3^)^16^.

### Tissue collection for tissue weight measurement and histological analysis

Testis, perigonadal white adipose tissue, epididymis and seminal vesicles were collected and weighted from animals euthanized by cervical dislocation under isoflurane anesthesia. Testis destined to H&E staining were fixed immediately in 4% paraformaldehyde overnight at 4°C, then dehydrated in gradient ethanol, made transparent in xylene, and embedded in paraffin. Embedded tissues were cut into 5 μm sections, which were deparaffinized and rehydrated for hematoxylin and eosin (H&E) staining.

### Fertility evaluation

#### F1 Male fertility with non-synchronized females

Adult F1-CD and F1-LPD male mice were caged with females from the animal supplier (Janvier Labs) at a 1:2 male to female sex ratio for 5 days. Plugs were checked every day and plugged females were removed and housed individually until parturition. The number of total females, plugged females, litters and offspring was counted to calculate the frequency of copulatory plug (FCP) and frequency of conception (FC).

#### F1 Female fertility

Adult F1-CD and F1-LPD female mice were caged with males from the animal supplier (Janvier Labs) at a 1:2 male to female sex ratio for 5 days. Plugs were checked every day and plugged females were removed and housed individually until parturition. The number of total females, plugged females, litters and offspring was counted to calculate the frequency of copulatory plug (FCP) and frequency of conception (FC).

#### F1 Male fertility with synchronized/superovulated females

Females from the animal supplier (Janvier Labs) were synchronized and superovulated using intraperitoneal injection of 5 units of pregnant mare serum gonadotrophin (PMSG) at 3pm on day 1, followed by 5 units of human chorionic gonadotropin (hCG) 48 h later (at 3pm on day3). Ovulation occurs 12h following the hCG injection (at 3am on day4). Adult F1-CD and F1-LPD male mice were caged with synchronized/superovulated females at a 1:2 male to female sex ratio for 15 hours either the night from day 1 to day 2 (from 6pm day1 to 9am day2, i.e. from 42h to 57h before ovulation) or the night from day 3 to day 4 (from 6pm day3 to 9am day4, i.e. from 9h before ovulation to 6h after ovulation). Plugs were checked the next morning and plugged females were removed and housed individually until sacrifice 14 days later.

### Sperm capacitation analysis

#### Capacitation media

The basic medium used throughout these studies for sperm preparation and in vitro capacitation was a modified Krebs Ringer medium (Whitten’s-HEPES buffer (WH) containing 100mM NaCl, 4.7mM KCl, 1.2mM KH2PO4, 1.2mM MgSO4, 5.5mM glucose, 1mM pyruvic acid, 4.8mM lactic acid, 20mM HEPES, pH 7.4). Capacitation medium contained 5mg/mL fatty-acid-free BSA (Sigma-Aldrich), 2mM CaCl2 and 20mM NaHCO 3 in WH.

#### Sperm collection

For all sperm analysis experiments, sperm were collected by pressure on cauda epididymis in 1.0mL of non-capacitating WH medium and then counted using a Malassez hemocytometer

#### Analysis of capacitation by Western blot

Sperm were incubated in capacitation medium at the final concentration of 15×106cells/mL during 0, 10, 30, 60, 90 or 120 min at 37°C, 5% CO2. At each time point, 0.75×106 cells were collected, centrifuged at 500g for five minutes, washed with 300μL of phosphate buffered saline (PBS), centrifuged at 500g for five minutes and then resuspended in Laemli buffer without β-Mercaptoethanol and boiled for five minutes. After a centrifugation at 7000g for five minutes, the supernatant was collected, boiled in the presence of 5% β-Mercaptoethanol (Sigma) for five minutes and then subjected to SDS-PAGE and transferred on nitrocellulose membrane (GE Healthcare #RPN3032D). Blots were blocked with Tris Buffered Saline (50mM Tris, 150mM NaCl) containing 0.1% v/v Tween 20 and 10% w/v low-fat dried milk. Membranes were probed overnight at 4°C with anti-phosphotyrosine antibody (1/1000, 05-321, clone 4G10, Merck Millipore) in blocking solution. After washing, membranes were incubated with the following secondary antibodies: anti-mouse horseradish peroxidase-conjugated (HRP) (Cell Signaling Technology # 7076S, 1/5000). Detection was performed using the Immobilon Western Chemiluminescent HRP substrate (Millipore Immobilon Crescendo Western HRP substrate # WBLUR0500) on Syngene G:BOX camera, and densitometric analyses were carried out using ImageJ software.

Normalization was performed by using anti-Tubulin antibody (Cell Signaling Technology # 2144S, 1/1000), in blocking solution. After washing, membranes were incubated with the following secondary antibodies: anti-rabbit horseradish peroxidase-conjugated (HRP) (Cell Signaling Technology # 7074S, 1/5000).

### Sperm motility analysis and In Vitro Fertilization (IVF)

#### Sperm collection

For all sperm analysis experiments, mouse sperm, obtained by manual trituration of caudae epididymis in 1mL M2 were allowed to swim out for 10 min at 37°C then counted using a Malassez hemocytometer. Sperm were then capacitated in M16 medium containing 2% fatty acid-free bovine serum albumin (BSA) at 37°C in a 5% CO2 incubator at a concentration of 2*10^6^ sperm cells /mL for either 1h (for motility analysis) or 12h (for IVF)

#### Sperm motility analysis

Before capacitation of after 1h capacitation, sperm suspensions were loaded on a 20-μm chamber slide (Leja Slide, Spectrum Technologies) and placed on a microscope stage at 37 °C. Sperm parameters (sperm count, progressive motility and hyperactivated sperm) were examined using a computer-assisted semen analysis (CASA) system (Hamilton Thorne Research, Beverly, MA).

#### In vitro fertilization and embryo development

Oocytes were collected from mature Balb/c females, synchronized by exposure to 5 units of pregnant mare serum gonadotropin (PMSG) and 5 units of human chorionic Gonadotropin (hCG). After 12h capacitation, sperm were washed with M16 then introduced into droplets containing >10 oocytes. Oocytes were incubated with 0.5×10^6^ capacitated sperm (100μL) (37°C, 5% CO2) in M16 medium, and unbound sperm were washed away after 4h incubation. Oocytes were washed and transferred to potassium simplex medium (KSOM) for culture to later stages. The percentage of the 2-cell embryos examined 24–30 h later was regarded as an indication of successful fertilization.

## Acknowledgments

We are grateful to the staff of the Installation Expérimentale de Nutrition (INRAE, site of Theix) for providing everyday care to the animals; Anne Terrisse-Lottier for her involvement in animals welfare; Joël Drevet (GReD, Université Clermont Auvergne) for helpful discussion and Rachel Guiton (GReD, Université Clermont Auvergne) for FACS analysis.

---

**Fig. S1.**
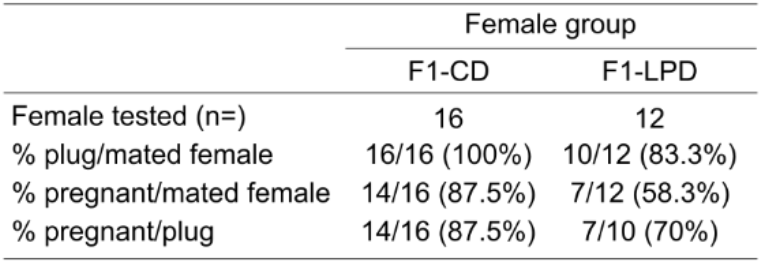
Fertility evaluation of non-synchronized F1-CD and F1-LPD Females when mated with Balb/c Males. F1-CD and F1-LPD Females were caged with Balb/c Males at Day 1 at a 1:2 male to female ratio for 4 days. Vaginal plugs were checked every day and plugged females were removed and isolated until delivery. For each group, the number of females tested, plugged females and pregnant females were indicated in the table. Fertility rates are calculated as the percentage of plugged females that are pregnant. Statistical significance was calculated using a Khi-squared test. ns = non-significant.

**Fig. S2.**
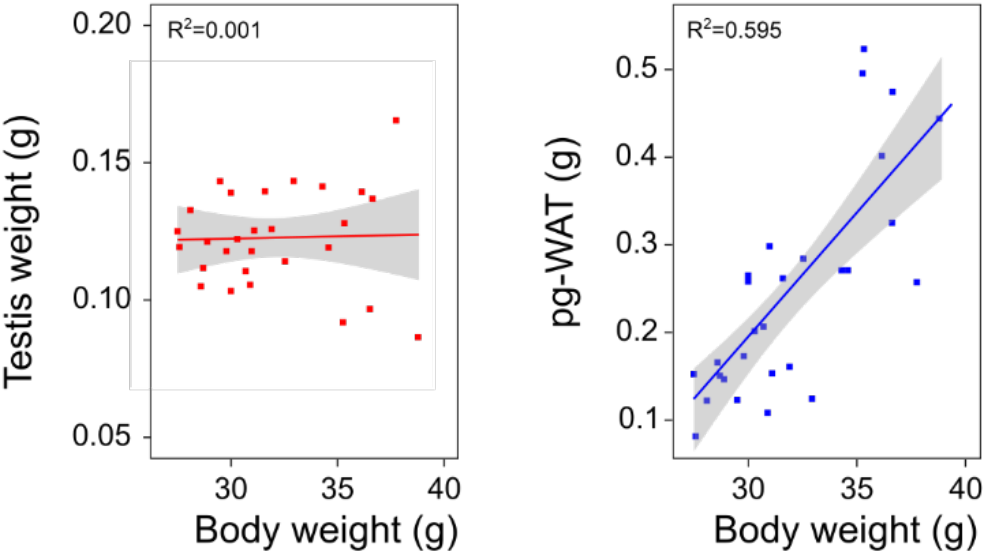
Correlation between Body weight and Testis weight or perigonadal-White Adipose Tissue weight (pg-WAT) in Balb/c males. The testis mass (in red) and perigonadal-WAT mass (in blue) of a large cohort of adult Balb/c males (n=28) were represented as a function of Body weight. R^2^ represents the R-squared of the linear regression model.

## References

1. Barker DJ, Intrauterine programming of adult disease. Mol Med Today 1:418–423 (1995).

2. Gingras V, Hivert M-F, Oken E, Early-Life Exposures and Risk of Diabetes Mellitus and Obesity. Curr Diab Rep 18:89 (2018).

3. Ong TP, Guest PC, Nutritional Programming Effects on Development of Metabolic Disorders in Later Life. Methods Mol Biol 1735:3–17 (2018).

4. Buckley AJ, Jaquiery AL, Harding JE, Nutritional programming of adult disease. Cell and Tissue Research 322:73–79 (2005).

5. Jousse C, Muranishi Y, Parry L, et al Perinatal protein malnutrition affects mitochondrial function in adult and results in a resistance to high fat diet-induced obesity. PLoS ONE 9:e104896 (2014).

6. Jousse C, Parry L, Lambert-Langlais S, et al, Perinatal undernutrition affects the methylation and expression of the leptin gene in adults: implication for the understanding of metabolic syndrome. FASEB J 25:3271–3278 (2011).

7. Mascarenhas MN, Flaxman SR, Boerma T, et al, National, Regional, and Global Trends in Infertility Prevalence Since 1990: A Systematic Analysis of 277 Health Surveys. PLoS Med 9:e1001356–12 (2012).

8. Kumar N, Singh AK, Trends of male factor infertility, an important cause of infertility: A review of literature. J Hum Reprod Sci 8:191–196 (2015).

9. Carlsen E, Giwercman A, Keiding N, Skakkebaek NE, Evidence for decreasing quality of semen during past 50 years. BMJ 305:609–613 (1992).

10. Wohlfahrt-Veje C, Main KM, Skakkebaek NE, Testicular dysgenesis syndrome: foetal origin of adult reproductive problems. Clinical Endocrinology 71:459–465 (2009).

11. Rae MT, Palassio S, Kyle CE, et al, Effect of maternal undernutrition during pregnancy on early ovarian development and subsequent follicular development in sheep fetuses. Reproduction 122:915–922 (2001).

12. Guzmán C, Cabrera R, Cárdenas M, et al, Protein restriction during fetal and neonatal development in the rat alters reproductive function and accelerates reproductive ageing in female progeny. J Physiol 572:97–108 (2006).

13. Zambrano E, Rodríguez-González GL, Guzmán C, et al, A maternal low protein diet during pregnancy and lactation in the rat impairs male reproductive development. J Physiol 563:275–284 (2005).

14. MacLeod DJ, Sharpe RM, Welsh M, et al, Androgen action in the masculinization programming window and development of male reproductive organs. International Journal of Andrology 33:279–287 (2010).

15. Scott HM, Hutchison GR, Jobling MS, et al, Relationship between Androgen Action in the “Male Programming Window,” Fetal Sertoli Cell Number, and Adult Testis Size in the Rat. Endocrinology 149:5280–5287 (2008).

16. Gallavan RH, Holson JF, Stump DG, et al, Interpreting the toxicologic significance of alterations in anogenital distance: potential for confounding effects of progeny body weights. Reprod Toxicol 13:383–390 (1999).

17. Wainstock T, Shoham-Vardi I, Sheiner E, Walfisch A, Fertility and anogenital distance in women. Reproductive Toxicology 73:345–349 (2017).

18. Clegg ED, Perreault SD, Klinefelter G, Assessment of male reproductive toxicology. Principles and Methods of Toxicology 1263–1299 (2001).

19. Olaso R, Habert R, Genetic and cellular analysis of male germ cell development. J Androl 21:497–511 (2000).

20. Gomendio M, Roldan ER, Coevolution between male ejaculates and female reproductive biology in eutherian mammals. Proc Biol Sci 252:7–12 (1993).

21. Hanson JL, Hurley LM, Female Presence and Estrous State Influence Mouse Ultrasonic Courtship Vocalizations. PLoS ONE 7:e40782–11 (2012).

22. Shukla KK, Mahdi AA, Rajender S, Ion Channels in Sperm Physiology and Male Fertility and Infertility. J Androl 33:777–788 (2012).

23. Santi CM, Orta G, Salkoff L, et al, Chapter Fourteen - K+ and Cl-Channels and Transporters in Sperm Function. In: Wassarman PM (ed) Gametogenesis Academic Press, pp 385–421 (2013)

24. Austin CR, The “Capacitation” of the Mammalian Sperm. Nature 170:326–326 (1952).

25. Travis AJ, Kopf GS, The role of cholesterol efflux in regulating the fertilization potential of mammalian spermatozoa. J Clin Invest 110:731–736 (2002).

26. Aitken RJ, Reactive oxygen species as mediators of sperm capacitation and pathological damage. Mol Reprod Dev 84:1039–1052 (2017).

27. Storey BT, Biochemistry of the induction and prevention of lipoperoxidative damage in human spermatozoa. Mol Hum Reprod 3:203–213 (1997).

28. Ecroyd HW, Jones RC, Aitken RJ, Endogenous redox activity in mouse spermatozoa and its role in regulating the tyrosine phosphorylation events associated with sperm capacitation. Biol Reprod 69:347–354 (2003).

29. Aitken RJ, Gibb Z, Mitchell LA, et al, Sperm Motility Is Lost In Vitro as a Consequence of Mitochondrial Free Radical Production and the Generation of Electrophilic Aldehydes but Can Be Significantly Rescued by the Presence of Nucleophilic Thiols1. Biol Reprod 87:367–11 (2012).

30. Nowicka-Bauer K, Nixon B, Molecular Changes Induced by Oxidative Stress that Impair Human Sperm Motility. Antioxidants 9:134–22 (2020).

31. Léonhardt M, Lesage J, Croix D, et al, Effects of perinatal maternal food restriction on pituitary-gonadal axis and plasma leptin level in rat pup at birth and weaning and on timing of puberty. Biol Reprod 68:390–400 (2003).

32. Aquila S, Rago V, Guido C, et al, Leptin and leptin receptor in pig spermatozoa: evidence of their involvement in sperm capacitation and survival. Reproduction 136:23–32 (2008).

33. Buffone MG, Verstraeten SV, Calamera JC, Doncel GF, High Cholesterol Content and Decreased Membrane Fluidity in Human Spermatozoa Are Associated With Protein Tyrosine Phosphorylation and Functional Deficiencies. J Androl 30:552–558 (2009).

34. Buffone MG, Human sperm subpopulations: relationship between functional quality and protein tyrosine phosphorylation. Hum Reprod 19:139–146 (2004).

35. Kothandaraman N, Agarwal A, Abu-Elmagd M, Al-Qahtani MH, Pathogenic landscape of idiopathic male infertility: new insight towards its regulatory networks. npj Genomic Medicine 1–9 (2016).

